# Flexible annotation atlas of the mouse brain: combining and dividing brain structures of the Allen Brain Atlas while maintaining anatomical hierarchy

**DOI:** 10.1101/2020.02.17.953547

**Authors:** Norio Takata, Nobuhiko Sato, Yuji Komaki, Hideyuki Okano, Kenji F. Tanaka

## Abstract

A brain atlas is necessary for analyzing structure and function in neuroimaging research. Although various annotation volumes (AVs) for the mouse brain have been proposed, it is common in magnetic resonance imaging (MRI) of the mouse brain that regions-of-interest (ROIs) for brain structures (nodes) are created arbitrarily according to each researcher’s necessity, leading to inconsistent ROIs among studies. One reason for such a situation is the fact that earlier AVs were fixed, *i.e.* combination and division of nodes were not implemented. This report presents a pipeline for constructing a flexible annotation atlas (FAA) of the mouse brain by leveraging public resources of the Allen Institute for Brain Science on brain structure, gene expression, and axonal projection. A mere two-step procedure with user-specified, text-based information and Python codes constructs FAA with nodes which can be combined or divided objectively while maintaining anatomical hierarchy of brain structures. Four FAAs with total node count of 4, 101, 866, and 1,381 were demonstrated. Unique characteristics of FAA realized analysis of resting-state functional connectivity (FC) *across* the anatomical hierarchy and *among* cortical layers, which were thin but large brain structures. FAA can improve the consistency of whole brain ROI definition among laboratories by fulfilling various requests from researchers with its flexibility and reproducibility.

**Highlights:**

– A flexible annotation atlas (FAA) for the mouse brain is proposed.
– FAA is expected to improve whole brain ROI-definition consistency among laboratories.
– The ROI can be combined or divided objectively while maintaining anatomical hierarchy.
– FAA realizes functional connectivity analysis *across* the anatomical hierarchy.
– Codes for FAA reconstruction is available at https://github.com/ntakata/flexible-annotation-atlas
– Datasets for resting-state fMRI in awake mice are available at https://openneuro.org/datasets/ds002551

## 1. Introduction

A standard brain atlas plays a key role in magnetic resonance imaging (MRI) for investigating anatomical and functional architecture of the brain (Aggarwal et al., 2011; Hess et al., 2018; Van Essen, 2002). Various three-dimensional annotation volumes (AVs) for the mouse brain have been proposed with various spatial resolution (32–156 μm), numbers of animals for averaging (4–27 animals), and numbers of segmented structures (5–70, and 707 regions-of-interest, ROIs) (Dorr et al., 2008; Johnson et al., 2010; Kovačević et al., 2005; Ma et al., 2008, 2005; Nie et al., 2019; Ullmann et al., 2013; Watson et al., 2017). Despite these excellent annotation atlases, it is still common in MRI studies of the mouse brain that ROIs for brain structures are prepared with arbitrary boundaries and locations according to each researcher’s necessities. For instance, a spherical ROI was used for the hippocampus and the cortex, which nevertheless did not reflect an actual spatial distribution (Takata et al., 2018, 2015). Such a situation might result from the fact that earlier AVs were fixed, *i.e.* combinations and divisions of ROIs for brain structures were not implemented, leading to inconsistent ROIs among laboratories for MRI in the mouse brain.

The Allen Institute for Brain Science (AIBS) provides resources related to the brain of C57BL/6J mouse (Sunkin et al., 2013). After two earlier editions in 2008 and 2011, AIBS provided a complete version for a common coordinate framework ver. 3 (CCFv3) for the mouse brain in 2015 (Allen Institute for Brain Science, 2017; Oh et al., 2014). Spatial coordinates of three-dimensional volume data at AIBS adopt CCFv3, enabling integration of multimodal resources by AIBS on 1) an anatomical template of the mouse brain (AT), 2) annotation volume (AV), 3) gene expression, and 4) axonal fiber projection. An increasing number of studies have used the Allen mouse brain atlas for analyzing MRI data (Grandjean et al., 2017b; Rubinov et al., 2015). Still, reconstruction of ROIs used in these studies is difficult because the original AVs by AIBS are often modified according to each researcher’s necessity without instruction for ROI reconstruction. Indeed, the original AV by AIBS *per se* is not perfectly suited for MRI analysis because a considerable number of brain structures defined in it are too small or large for MRI analysis.

This report presents a pipeline to construct a flexible annotation atlas (FAA) for the mouse brain, for which brain structures can be combined or divided objectively using resources by AIBS on gene expression and fiber projection. Total ROI counts of FAA can vary from 1 to more than 1,000. Construction of FAA takes just two steps, combining and dividing brain structures, using user-specified, text-based information and Python codes. FAA enables analysis of resting-state functional connectivity (FC) across the anatomical hierarchy because FAA retains information related to the hierarchy. Furthermore, cortical layer-specific FC was examined using FAA that had thin but large ROIs for cortical layers. FAA would present one more choice for a consistent ROI definition for the mouse brain among laboratories by satisfying various requests from researchers with its flexibility, simplicity, and reproducibility.

## 2. Materials and Methods

### 2.1 Ethics statement

All animal experiments were conducted in accordance with the National Institutes of Health Guide for Care and Use of Laboratory Animals (NIH Publications No. 8023) and were approved by the Animal Research Committee of Keio University (approval number: 12034-(5)) and CIEA (16062A and 16048R).

### 2.2 Public resources on the mouse brain by the Allen Institute for Brain Science

Online public resources by AIBS used for construction of FAA included a text file for annotation ontology (AO; Fig. 1a right), and four volume data: 1) anatomical template of the mouse brain (AT; Fig. 1a left), 2) annotation volume (AV; Fig. 1a middle), 3) gene expression data (Fig. 3c Upper panels), and 4) axonal fiber projection data (Fig. 3d Upper panels). Their download links are presented in Supplementary Table 1.

**Fig. 1.**
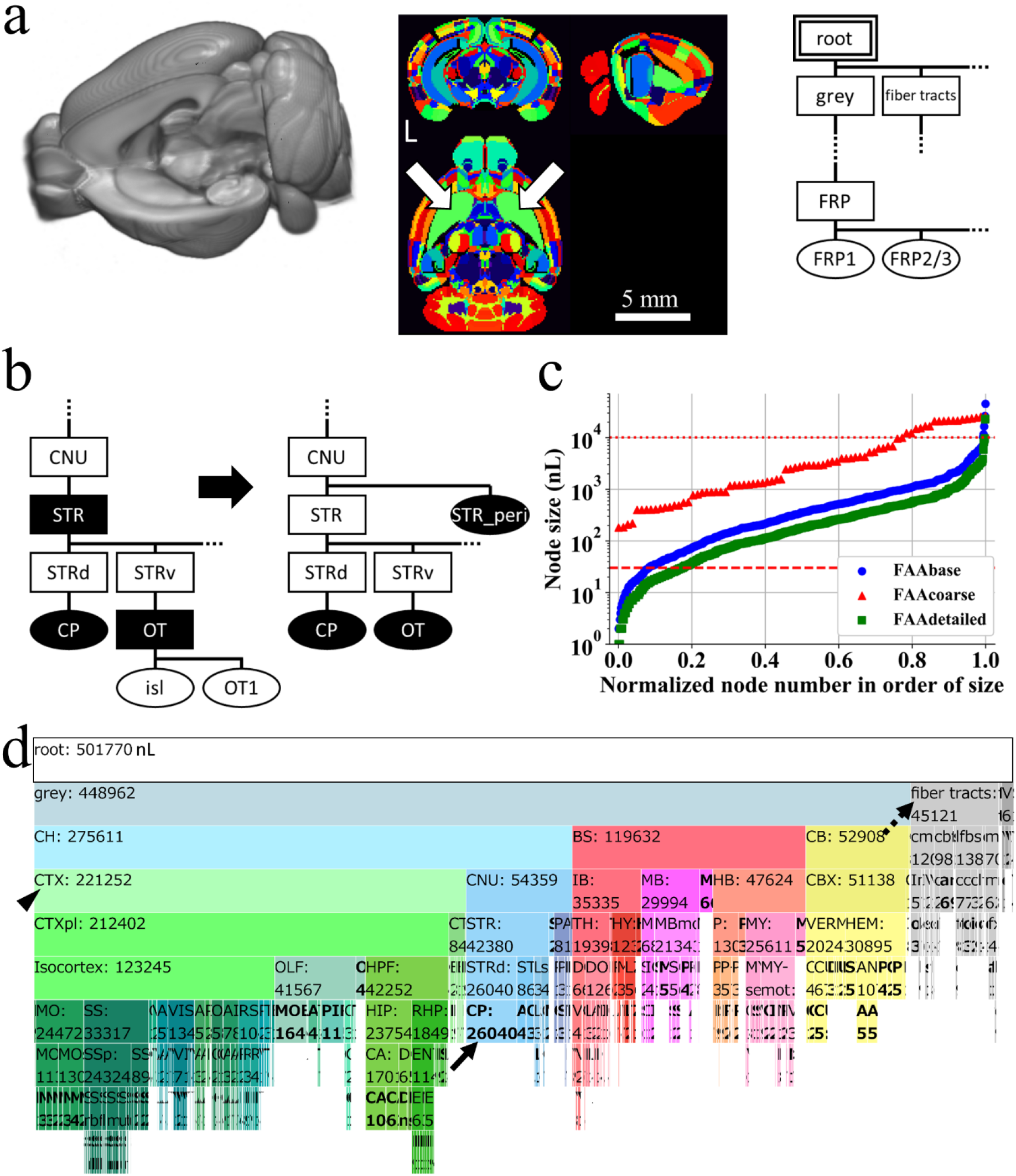
Preprocessing of resources by Allen Institute for Brain Science. **(a)** Online public resources at AIBS for anatomical template (AT, left), annotation volume (AV, middle), and anatomical ontology (AO, right) of the mouse brain. AT is shown with its left hemisphere cropped to display its subcortical structures. AV is shown in NIH color scheme, with each voxel assigned an integer value for an ID of a brain structure. This original AV is single-sided: homotopic brain structures on the left and right sides of the brain have the same ID and constitute a single node. The same brain structure (ID) is shown in the same color, *e.g.* arrows indicate a single node in green color, caudoputamen (CP). L denotes the left side of the brain. AO is a JSON-formatted text file shown here as a rooted tree with each brain structure regarded as a *node* in a graph theoretical term, demonstrating its anatomical hierarchy. A double line rectangle shows a root node of a rooted tree graph. A parent node is one-level higher than its child nodes in the anatomical hierarchy. For example, a node “root” is a parent node of its child nodes “grey” and “fiber tracts”. Similarly, a node “FRP” is a parent node of its child nodes “FRP1” and “FRP2/3”. Nodes “grey and fiber tracts” and “FRP1 and FRP2/3” respectively occupy the same level of the hierarchy. Ellipses and rectangles respectively denote leaf nodes, which do not have a child node, and inner nodes, which have at least one child node, with its acronym for a brain structure: grey, grey matter; FRP, Frontal pole in the cerebral cortex; FRP1 and 2/3, Layer 1 and 2/3 of FRP. **(b)** Preprocessing of the original AO (left) for eliminating *destructive* nodes to obtain a modified AO (designated as AObase; right). The original AV was updated to AVbase by reflecting AObase. A pair of AObase and AVbase was designated collectively as a flexible annotation atlas base (FAAbase). Filled and open nodes: nodes with and without corresponding voxels in AV. CNU, cerebral nuclei; STR, striatum; STRd and STRv, dorsal and ventral region of STR; OT, olfactory tubercle; isl, islands of Calleja; OT1, molecular layer of OT. **(c)** Distribution of size (voxel count in AVbase) of leaf nodes in FAAbase (blue plots). Too small or too large nodes for fMRI analysis are evident in FAAbase. Specifically, 6.7% of all leaf nodes in FAAbase were smaller than a single voxel in fMRI (30 nL, dashed line). Four nodes were extremely large (> 10,000 nL, dotted line). Red and green plots are for FAAcoarse and FAAdetailed (see Figs. 2 and 3). **(d)** This icicle plot presents the anatomical hierarchy of brain structures in FAAbase. Each node is shown with its acronym and size in units of nL. The width of each node represents its size. Root nodes are at the top; leaf nodes (shown in bold face) are at the bottom such as CP in blue (indicated by the arrow at the middle). Inner nodes are between a root node and leaf nodes, such as cerebral cortex (CTX) in pale green (indicated by the arrowhead at the left). Nodes for fiber traces and ventricular systems (VS) are included in FAAbase (indicated by the dashed arrow at the upper right). FAAbase is also single-sided. For nodes with collapsed letters, see the zoomable HTML version of this plot. The color code for the brain structure follows that of AIBS, and is applicable to Figs. 2b, 2d, 3e, 4a, 4b, 4d, and Supplementary Fig. 1a. CH, cerebrum; BS, brain stem; CB, cerebellum. For other acronyms, see the original AO file.

**Fig. 2.**
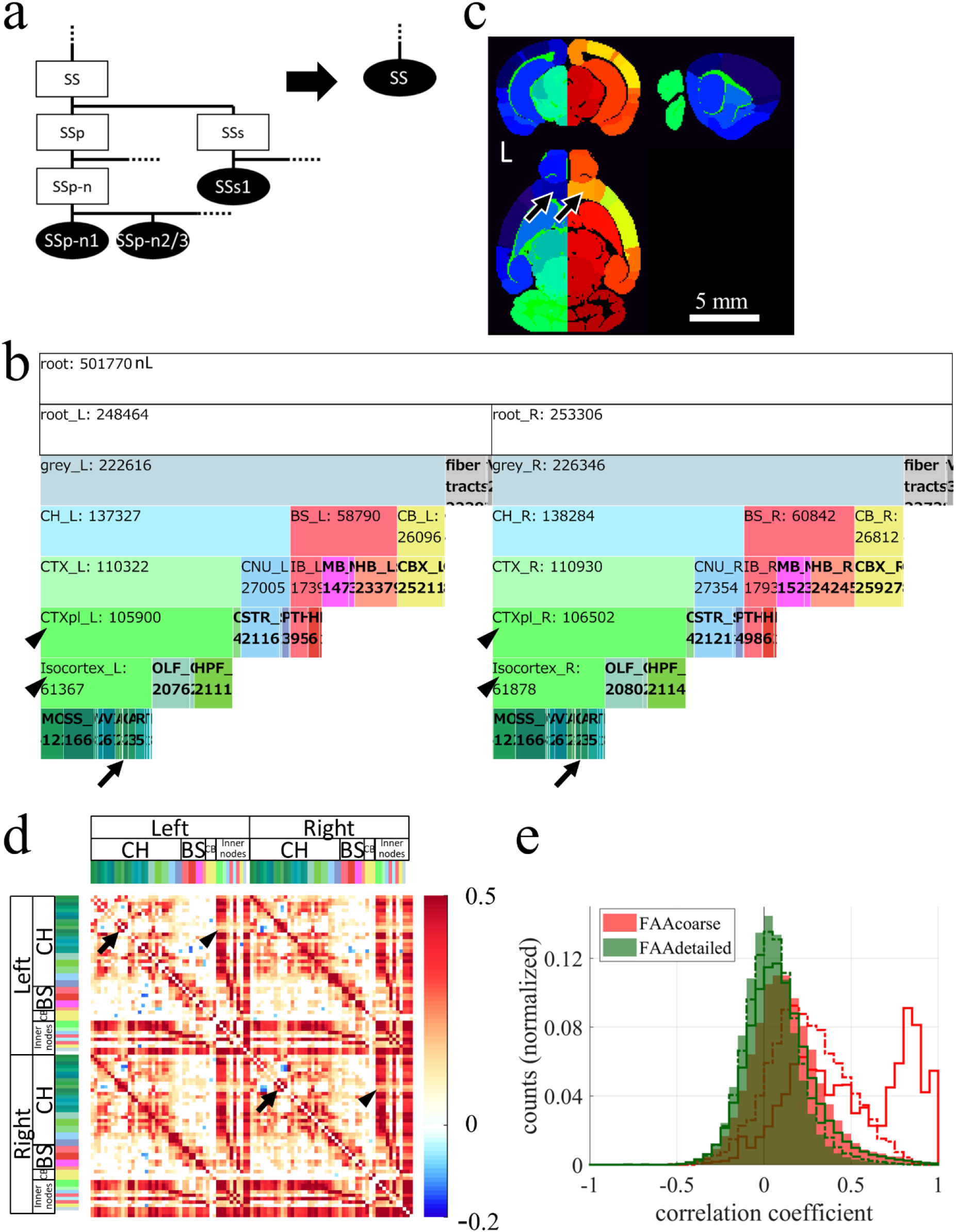
Combining nodes to create a larger node while maintaining anatomical hierarchy. **(a)** Example of combining nodes. Leaf nodes (SSp-n1, SSp-n2/3, and others) and inner nodes (SSp, SSs, and others), which were both descendant nodes of an *inner* node SS in AObase (left), were combined to create a *leaf* node SS, obtaining AOcoarse (right). AVbase was updated to AVcoarse by reflecting AOcoarse. A pair of AOcoarse and AVcoarse was designated collectively as FAAcoarse. Combining all descendant nodes of SS was necessary to maintain the anatomical hierarchy. Ellipses and rectangles respectively denote leaf nodes and inner nodes. Filled and open nodes respectively represent nodes with and without corresponding voxels in AV: SS, somatosensory areas; SSp, primary SS; SSp-n, nose region of SSp; SSp-n1 and n2/3, layer 1 and layer 2/3 of SSp-n; SSs, supplemental SS; SSs1, layer 1 of SSs. **(b)** An icicle plot of anatomical hierarchy of FAAcoarse. A root node at the top has child nodes, root_L and root_R, indicating that FAAcoarse is double-sided, *i.e.* different IDs are assigned to homotopic nodes in the left and right side of the brain. Zoomable HTML version of this plot is available. Arrows point leaf nodes in the prefrontal cortex (PFC), *i.e.* prelimbic area (PL), infralimbic area (ILA), and orbital area (ORB) on both sides of the brain. Arrowheads indicate ancestor nodes for the PFC, *i.e.* cortical plate (CTXpl) and isocortex. **(c)** Sectional images of AVcoarse shown in NIH color scheme. Nodes in the left and right side of the brain have, respectively, bluish and reddish colors reflecting that nodes in the right hemisphere were assigned larger IDs. For example, IDs in FAAcoarse for nodes ORB (arrows) in the PFC on the left and right sides of the brain are, respectively, 18 and 68. Larger node size is evident in this AVcoarse than AVbase (Fig. 1a middle). L denotes the left side of the brain. **(d)** Resting-state FC matrix calculated using FAAcoarse. FAAcoarse realized FC analysis *across* the anatomical hierarchy because FAAcoarse retains hierarchy-related information. *Inner* nodes such as the cerebral cortex (CTX, pale green in color codes) and its descendant nodes typically had higher FC than leaf nodes. This higher FC is evident in vertical and horizontal red bands in the FC matrix, demonstrating high FCs between *inner* nodes and between an inner node and a leaf node in the cerebral cortex. One exception was found in the PFC. Whereas high FC was found between *leaf* nodes in the right and left PFC (arrows) such as PL, ILA, and ORB, low FC was observed between the leaf nodes and inner nodes such as CTXpl and Isocortex (arrowheads). Only nodes for brain parenchyma were used for FC analysis, excluding nodes for fiber tracts and ventricles. The color bar at the right shows Pearson’s correlation coefficient. Color codes at the top and left represent brain structures. For inner nodes, color codes match those in Fig. 2b. **(e)** Histogram of correlation coefficients between leaf nodes (filled histogram), between a leaf node and an inner node (dotted line), and between inner nodes (solid line) in the FC matrix by FAAcoarse (red color). A shift to the right of solid and dotted lines compared to a filled histogram confirms higher FCs for inner nodes. Especially strong FCs around 0.8–1.0 were observed between inner nodes “CH and CTX” and “CTXpl and Isocortex”. The green histogram is for FAAdetailed (Fig. 4a).

**Fig. 3.**
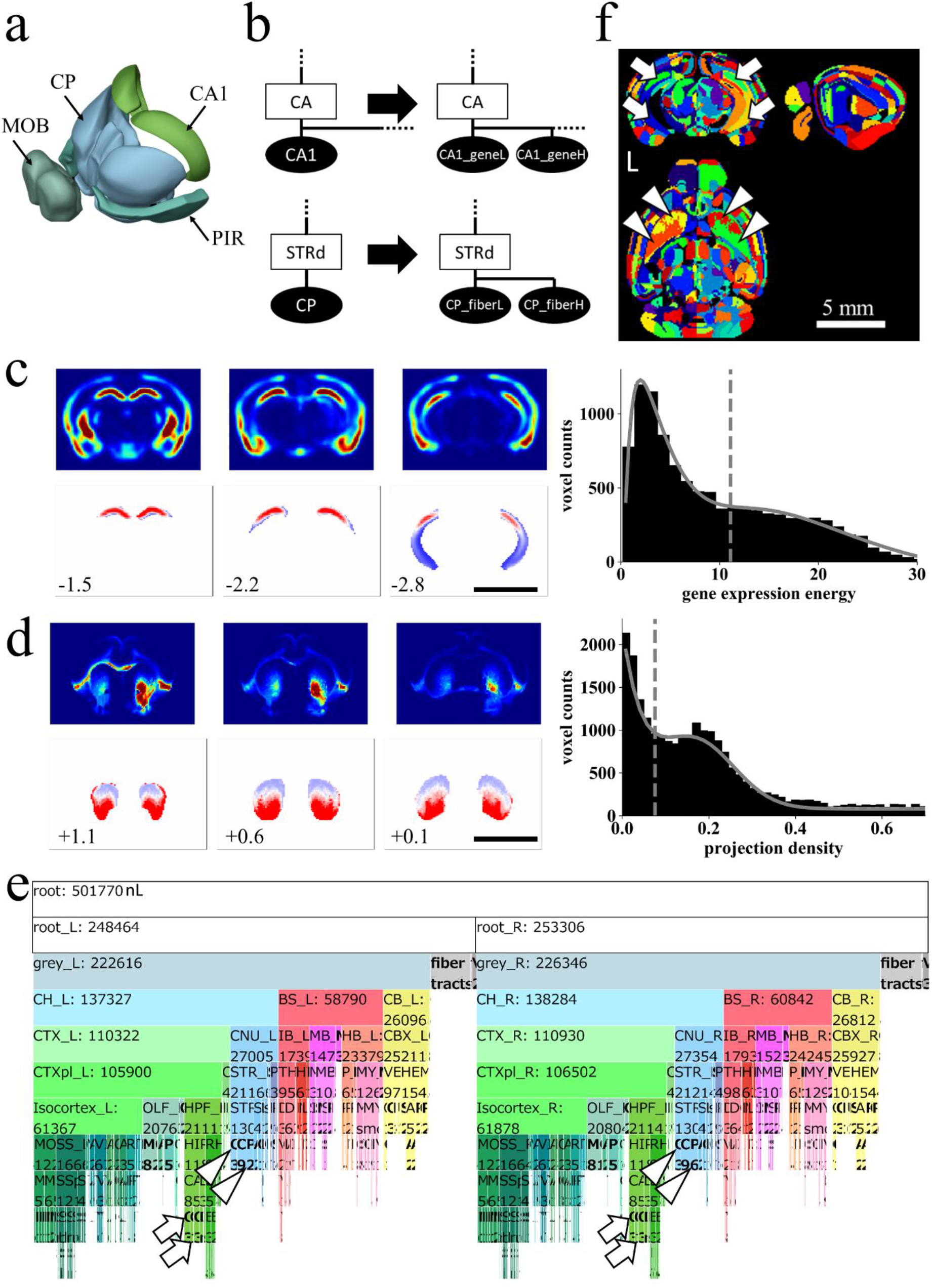
Dividing nodes using resources by AIBS on gene expression and fiber projection. **(a)** The four largest nodes in the original AV: CP, Caudoputamen, 26,040 nL; MOB, Main olfactory bulb, 16,406 nL; PIR, Piriform area, 11,591 nL; CA1, Field CA1 in the hippocampus, 10,278 nL. **(b)** Example of dividing nodes. Leaf nodes in AObase (left) were divided to new leaf nodes using resources at AIBS on gene expression or axonal fiber innervation, obtaining AOdetailed (right). Ellipses and rectangles respectively denote leaf nodes and inner nodes. Filled and open nodes signify nodes with and without corresponding voxels in AV: CA, Ammon’s horn. **(c, d)** Dividing leaf nodes CA1 and CP based respectively on a gene-expression energy and axonal projection density. **Upper panels:** Coronal sections of volume data obtained at AIBS for gene-expression energy of a gene *Wfs-1* (**c**) or axonal projection-density from the insula cortex to CP (**d**). **Right:** Histograms of the gene-expression energy within a node CA1 fitted with two Gaussian curves (**c**) and of the projection-density within a node CP fitted with a Gaussian curve and a Poisson distribution curve (**d**). Dashed line: an inflection point of the fitted curve used as a threshold. **Lower panels:** Divided nodes. Leaf nodes CA1 and CP were divided, respectively, to two by the thresholds, resulting in nodes “CA1_geneH (red) and CA1_geneL (blue)” and “CP_fiberH (red) and CP_fiberL (blue)”. Axonal projection data were flipped horizontally to obtain symmetrical projection data before thresholding. Values at the lower left represent the anterior–posterior (AP) distance from bregma in millimeters. Scale bar: c, d, and f, 5 mm. **(e)** An icicle plot of anatomical hierarchy of FAAdetailed. Arrows indicate divided leaf nodes based on gene expression in the left (CA1_geneL_L and CA1_geneH_L) and the right (CA1_geneL_R and CA1_geneH_R) side of the brain. Arrowheads show divided leaf nodes based on fiber projection density in the left (CP_fiberL_L and CP_fiberH_L) and the right (CP_fiberL_R and CP_fiberH_R) side of the brain. A zoomable HTML version of this plot is available. **(f)** Sectional images of AVdetailed shown in random color scheme. Divided nodes for CA1 (arrows) and CP (arrowheads) on both sides of the brain are shown. Specifically, arrows indicate nodes “CA1_geneL_L” (ID 322, blue) and “CA1_geneH_L” (ID 323, green) in the left brain, and nodes “CA1_geneL_R” (ID 1006, yellow) and “CA1_geneH_R” (ID 1007, green) in the right brain. Similarly, arrowheads indicate nodes “CP_fiberL_L” (ID 372, orange) and “CP_fiberH_L” (ID 373, lime) in the left brain, and nodes “CP_fiberL_R” (ID 1056, green) and “CP_fiberH_R” (ID 1057, red) in the right brain.

Of those five resources, the first, AO, is a text file in JSON format with 17,549 lines that define the anatomical hierarchy of brain structures, which includes 1) a structure name, 2) acronym, 3) ID for a brain structure, 4) a parent structure’s ID, and others. For this study, each brain structure was regarded as a *node* in a graph theoretical term. A node at the top of an anatomical hierarchy in AO is a *root* node defined as an ancestor of every node (Fig. 1a right). Every node has a unique parent node except the root. An *inner* node is a node with at least one child node, *e.g.* a node striatum (STR) in Fig. 1b that has child nodes striatum dorsal region (STRd) and striatum ventral region (STRv). A *leaf* node is a node without a child node, *e.g.* a node caudoputamen (CP) in Fig. 1b. The original AO defines 1,327 brain structures, including 637 in the cerebrum, 375 in the brain stem, and 87 in the cerebellum, among which 197 are inner nodes and 1,130 are leaf nodes. The anatomical hierarchy defined in AO was visualized as rooted tree graphs (*e.g.* Fig. 1a right), icicle plots (*e.g.* Fig. 1d), or a compound spring embedder (CoSE) layout (Fig. 4d).

**Fig. 4.**
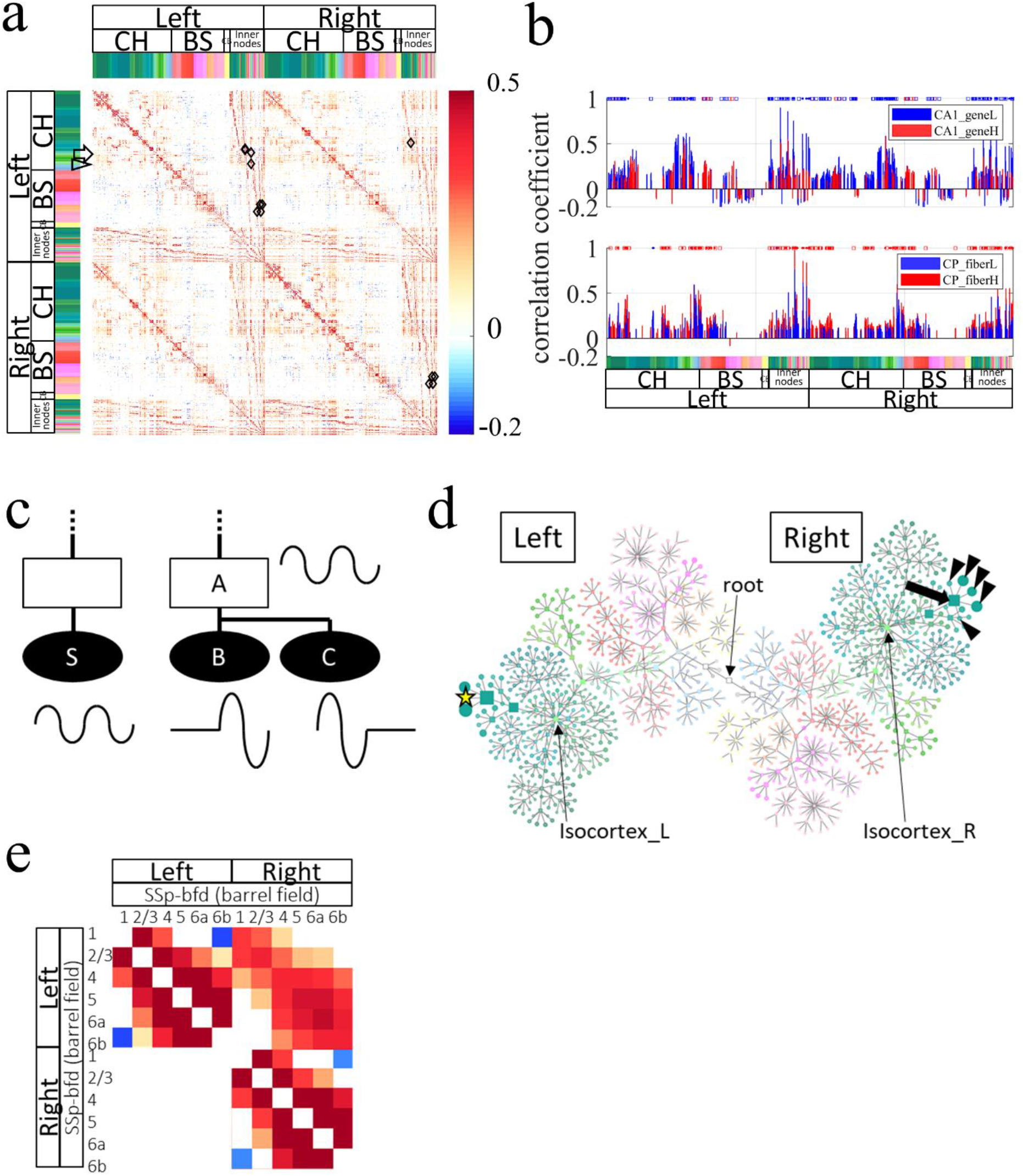
Resting-state FC across the anatomical hierarchy and between cortical layers. **(a)** Resting-state FC matrix calculated with FAAdetailed. FAAdetailed realized examination of FC modulation by an inner node because FAAdetailed retains information of the hierarchy. The arrow and arrowhead at the left respectively indicate rows of the FC matrix for divided nodes in CA1 and CP (see Fig. 4b). Diamonds show 13 pairs of a source leaf node and an inner node that satisfies the situation described below in (c). Non-significant correlation coefficients in the FC matrix were set to 0. **(b)** Bar graphs comparing FCs from the divided nodes in CA1 or CP in the left hemisphere. **Upper panel:** Distribution of mean correlation coefficients from a node CA1_geneL (ventral CA1, blue bars) and a node CA1_geneH (dorsal CA1, red bars) in the FC matrix. Colors of markers at the top show that FC from the ventral CA1 was significantly higher (blue) or lower (red) than FC from the dorsal CA1. Dominance of blue markers over red ones suggests stronger FCs from the ventral CA1 than that from the dorsal CA1. Markers in dots or squares respectively signify that FCs from both the ventral and dorsal CA1 were significant and that FC from only either one was significant in the FC matrix. Instances of more numerous squares over dots suggest distinct patterns of FCs from the ventral and dorsal CA1. **Lower panel:** Distribution of mean correlation coefficients from a node CP_fiberL (dorsal CP, blue bars) and a node CP_fiberH (ventral CP, red bars) in the FC matrix. Dominance of red markers over blue ones at the top suggests stronger FC from the ventral CP than from the dorsal CP. Instances of more numerous squares over dots suggest distinct patterns of FCs from the ventral and dorsal CP. Color code at the bottom represents brain structures. **(c)** Conceptual illustration for activity modulation by a larger unit in the brain, *i.e.* an *inner* node, that occupies a higher position in the anatomical hierarchy than *leaf* nodes occupy. Earlier studies have analyzed only FCs between leaf nodes, *e.g.* FC from a source leaf node S to leaf nodes (B and C). Presuming that FC from a source leaf node S to an *inner* node A is larger than FC to descendant *leaf* nodes (B and C) of the inner node A, which is not an ancestor node of the source node S, then this situation suggests a mechanism to modulate FC in larger units in the brain, *i.e.* an inner node A. Waves at each node represent the resting-state activity (BOLD signal). Ellipses and rectangles respectively denote the leaf node and inner node. Filled and open nodes signify nodes with and without corresponding voxels in AV. **(d)** Visualization of FCs with anatomical hierarchy from one source leaf node that satisfied the situation. A source leaf node (RSPv2/3_L, a yellow star at left) in the left brain was found to have higher FC to an *inner* node (RSPv_R, an arrow at right) in the right brain than that to descendant five leaf nodes (arrowheads at right) of the inner node. However, the difference of FC was not always significant. Similar results were obtained for all source leaf nodes. These results do not support the idea of FC modulation by a larger unit in the brain, at least during resting-state static FC of awake mice. Circles and squares respectively represent leaf nodes and inner nodes. The node size reflects the strength of FC. Color code is for brain structures. Left and Right respectively denote groups of nodes on the left and right sides of the brain. Thin arrows indicate nodes for a root, isocortex in the left brain (Isocortex_L), and isocortex in the right brain (Isocortex_R): RSPv2/3_L, layer 2/3 of retrosplenial area, ventral part in the left brain; RSPv_R, retrosplenial area, ventral part in the right brain. **(e)** Analysis of cortical-layer-specific FC that was enabled by FAAdetailed with thin but large nodes for cortical layers. This is an expanded view of the FC matrix (Fig. 4a) for the primary somatosensory area for barrel cortex (SSp-bfd). Significant difference in FC within *vs.* between supra-granular and infra-granular layers was found in the most cortical regions. Specifically, FC between L1 and L2/3 or between L5 and L6a was significantly larger than FC between L2/3 and L6a.

The second of those five resources, AT, is a three-dimensional average volume of the mouse brain constructed with serial two-photon tomography using 1,675 specimens. The third, AV, is an annotated brain volume, the voxels of which are assigned to an integer value representing ID for a brain structure that is defined in AO. The original AV has 670 unique integer values: from 0, which represents outside of the brain, to 614,454,277, which represents the supraoculomotor periaqueductal gray. The fourth, gene expression volume, was prepared with genome-wide *in situ* hybridization (ISH) image for approximately 20,000 genes in adult mice (Allen Institute for Brain Science, 2014). Finally, the fifth, the axonal fiber projection volume, was constructed with serial two-photon tomography using viral tracers for 2,995 axonal pathways (Allen Institute for Brain Science, 2018; Kuan et al., 2015).

Actually, AIBS provides several three-dimensional volume data for AT, AV, and axonal projection density with different spatial resolutions (10– 100 μm isovoxel; Supplementary Table 1). For this study, spatial resolution of 100 μm isovoxel was used, although construction of FAA was compatible with any resolution, because 1) the spatial resolution best matched that of our functional MRI (fMRI, 200 × 200 × 750 μm^3^), 2) the total count of unique IDs in AVs with higher spatial resolution did not increase to a considerable degree (669, 670, 671, and 672 unique IDs in AVs with spatial resolution of 100, 50, 25, and 10 μm isovoxel, respectively), and 3) a single voxel of the volume data was exactly 1 nL.

### 2.3 Construction pipeline for a flexible annotation atlas of the mouse brain

FAA consists of a JSON-formatted text file (anatomical ontology, AO) and a three-dimensional volume file of the mouse brain (annotation volume, AV). First, the original files AO and AV by AIBS were preprocessed using Python codes without user inputs for eliminating *destructive* nodes in the original files to obtain AObase and AVbase (Fig. 1b; see 3.1 for *destructive*). The pair of AObase and AVbase was designated collectively as FAAbase. Numerous patterns of FAA can then be constructed using two steps with user-specified text-based information: 1) “combining leaf nodes” of FAAbase to produce a new leaf node with larger size, *i.e.* voxel counts in AV (Fig. 2), and 2) “dividing a leaf node” of FAAbase using resources at AIBS on gene expression and axonal projection (Fig. 3).

These steps were performed objectively using Python codes while maintaining 1) anatomical hierarchy defined in AO, and 2) consistency between AO and AV. Details of these procedures are summarized in Supplementary Tables 2–4. Four FAAs with different total ROI counts were constructed for the present study to demonstrate its flexibility: FAAbase (total node count: 866, leaf node count: 669; Fig. 1d), FAAsegment (4, 3; Supplementary Fig. 1), FAAcoarse (101, 80; Figs. 2b, 2c), and FAAdetailed (1381, 1097; Figs. 3e, 3f).

Specifically, the original AO was preprocessed to obtain AObase without destructive nodes, which means that leaf nodes and inner nodes in AObase do and do not have corresponding voxels in the original AV, respectively (Fig. 1b right). The original AV was updated accordingly to obtain AVbase. Four steps achieved this: 1) eliminating *leaf* nodes in the original AO that did *not* have corresponding voxels in the original AV, *e.g.* leaf nodes islands of Calleja (isl) and olfactory tubercle, molecular layer (OT1) in Fig. 1b; 2) dividing *inner* nodes in the original AO that *had* corresponding voxels in the original AV into two: a leaf node and an inner node respectively with and without corresponding voxels in the AV, *e.g.* an inner node STR in Fig. 1b; 3) updating IDs in the original AV by reflecting AObase to obtain AVbase; and 4) appending node size information in AVbase to AObase.

The “combining leaf nodes” procedure was performed by editing the AObase text file (Fig. 2a). This procedure was left as manual rather than automated by program codes because simple criteria such as thresholding by node size turned out to be insufficient to achieve flexibility for combining nodes. Indeed, the desirable threshold for a node size might be different, for example, in the cortex and the brainstem. Moreover, combining nodes is straightforward and easily accomplished using a text editor merely by deleting values of a key “children” for an inner node in the JSON-formatted text file, AObase. To facilitate this step further, a zoomable visualization of anatomical hierarchy of the brain structures in AO is provided as an HTML file using D3.js (https://d3js.org/) (*e.g.* Fig. 1d).

The procedure “dividing a leaf node” was performed with Python codes and user-specified text-based information related to 1) IDs for brain structures to be divided, 2) brain structure acronyms for source and target of axonal fiber innervation, and 3) experimental IDs found at a website of AIBS for gene expression (https://mouse.brain-map.org/) and axonal fiber projection (http://connectivity.brain-map.org/) (Fig. 3). Concretely, a node was divided to two using a threshold for 1) gene expression energy in the node that was defined as “gene expression intensity × expression density of ISH image for a gene” (Fig. 3c), or 2) fiber projection density from a source to the target-brain structure (Fig. 3d). The thresholds were defined objectively as an inflection point of fitted curves in histograms for gene expression energy or fiber projection density in a node (Fig. 3c or 3d right).

### 2.4 Resting-state fMRI acquisition in awake mice

Resting-state fMRI was performed in awake mice using a CryoProbe, as described previously (Matsubayashi et al., 2018; Takata et al., 2018; Yoshida et al., 2016). Briefly, acrylic head bar (3 × 3 × 27 mm^3^) was mounted using dental cement (Super-Bond C&B Sun Medical Co., Ltd., Shiga, Japan) along the sagittal suture of the exposed skull of seven male C57BL/6J mice anesthetized with 2–3% isoflurane. After recovery from surgery, mice were acclimated to a mock fMRI environment for 2 hr/day for at least 7 days before performing fMRI for awake mice. This standard procedure in *in vivo* two-photon imaging for awake mice achieves stable measurements of brain activity over several hours (Seibt et al., 2017; Takata et al., 2011) while avoiding confounding effects of anesthetics during fMRI because anesthesia is not necessary to place awake mice in an animal bed of MRI (Gao et al., 2017).

Structural and functional MRI was performed using a 7.0 Tesla MRI apparatus equipped with actively shielded gradients at 700 mT/m maximum strength (Biospec 70/16; Bruker BioSpin AG, Fällanden, Switzerland) with a cryogenically cooled 2-ch transmit/receive phased array surface coil (CryoProbe, Z120046; Bruker BioSpin AG, Fällanden, Switzerland) and a ParaVision 6.0.1 software interface (Bruker Biospin AG, Fällanden, Switzerland). Parameter tuning was performed before MRI acquisition: 1) manual tuning and matching of radiofrequency (RF) coils (wobble adjustment), 2) automatic adjustment of the resonance frequency (basic frequency), 3) calibrating the RF pulse power (reference power), 4) adjusting global linear shims (FID shim), 5) measuring B0 map, and 6) localized field-map shimming based on B0 map (MAPSHIM; repetition time [TR] = 20 ms; echo time [TE] = 1.520 ms, 5.325 ms; spatial resolution = 300 × 300 × 300 μm^3^; matrix = 64 × 64 × 64 voxels). After this tuning, T2-weighted structural images were acquired from 7 mice using a rapid acquisition process with a relaxation enhancement (RARE) sequence in coronal orientations (TR/TE = 6100/48 ms; spectral bandwidth [BW] = 5 kHz; RARE factor = 8; number of averages = 4; number of slices = 52; spatial resolution = 75 × 75 × 300 μm^3^). In all, 13 runs of T2* weighted fMRI were obtained using 7 mice with a gradient-echo echo-planar imaging (EPI) sequence (TR/TE = 1500/15 ms; N segments = 1; BW = 250 kHz; flip angle = 70°; FOV = 19.2 × 19.2 mm^2^; matrix = 64 × 64 × 18; number of slices = 18; slice thickness = 0.70 mm; slice gap = 0.05 mm; spatial resolution = 200 × 200 × 750 μm^3^; temporal resolution = 1.5 s; dummy scan = 0; number of repetitions = 400; scan time = 10 min). The fMRI scanning covered the whole brain including the olfactory bulb and the cerebellum.

### 2.5 Spatial preprocessing of MRI data

Spatial preprocessing of MRI data was performed using Advanced Normalization Tools (ANTs 2.1.0; http://stnava.github.io/ANTs/), Brain Extraction Tool (BET) (Smith, 2002), and SPM12 (7487, Welcome Trust Centre for Neuroimaging, London, UK). Concretely, structural MRI images in NIfTI format underwent the following: 1) N4 bias-field corrected by ANTs; 2) skull-stripped with a BET plugin in Multi-image Analysis GUI software (Mango v4.1 [1531]); 3) replicated with copying s-form code to q-form code about image orientation in a NIfTI header of the images because the BET procedure deleted q-form code; and 4) deformably registered to the original AT from AIBS in NIfTI format in RAS orientation by ANTs (antsRegistrationSyNQuick.sh). Similarly, time-series fMRI images were processed: 1) the first four volumes were taken away as dummy scans corresponding to the first 6 s of imaging (SPM file conversion); 2) images were realigned for head-motion correction (SPM-realign: Estimate & Reslice; Quality, 1; Separation, 3 mm; Smoothing, 5 mm; Register to mean; Interpolation with 2^nd^ degree B-spline); 3) images were corrected for slice-timing (SPM-slice timing); 4) images were reoriented to RAS orientation (SPM-check reg); 5) images were registered to the preprocessed structural image by ANTs using information related to rigid transformation of a mean fMRI image to the structural image (antsApplyTransforms); and finally 6) images were registered to the original AT from AIBS by ANTs using information related to transformation of the structural image to the AT (antsApplyTransforms). The final step (6) modulated spatial resolution of fMRI images to be 100 μm isovoxels, the same as the original volumes by AIBS. Spatial averaging of fMRI images using a Gaussian kernel was not used for this study because the isotropic characteristics of the kernel did not reflect an actual spatial distribution of brain structures, which might result in contamination of blood oxygenation level-dependent (BOLD) fMRI signals from other nodes, especially in a thin but large node such as a cortical layer (Fig. 4e). Instead, BOLD-fMRI signals were spatially averaged within each node defined in FAA to improve the signal-to-noise ratio (SNR).

### 2.6 Temporal preprocessing of fMRI data

Denoising of the fMRI time series was conducted using a functional connectivity toolbox (CONN 18.b). First, outlier scans were identified for scrubbing using artifact detection tools (ART). Concretely, fMRI volumes with spikes or framewise displacement greater than 50 μm that corresponds to 0.25 voxel of in-plane resolution, and/or with rotation greater than 1.15 degree were detected for usage as regressors. We intended to discard a whole run if confounding volumes exceeded 20% of all volumes, but this did not occur. Removal of confounding effects by linear regression was performed using 1) linear/quadratic trends within each functional run, 2) realignment-based subject motion, 3) noise signals in white matter and CSF, obtained with component-based noise correction method (CompCor), and 4) 0.01–0.1 Hz band-pass filtering. FAAsegment was substituted for a tissue probability map (TPM) based on image intensity of structural MRI (Hikishima et al., 2017; Meyer et al., 2017; Nie et al., 2019; Sawiak et al., 2009). FAAsegment was used for segmenting grey matter, white matter, and cerebrospinal fluid (CSF) in CONN analysis (CompCor) because mis-assignments of nodes in grey matter to that in white matter in a TPM were noticed, *e.g.* globus pallidus, external segment (GPe), ventral posterolateral nucleus of the thalamus (VPL), ventral posteromedial nucleus of the thalamus (VPM), pretectal region (PRT), retrosplenial area (RSP), and inferior colliculus (IC) (Hikishima et al., 2017). In addition, the original AT by AIBS is based on samples that are larger than commonly available TPMs (≤ 60 specimens). Finally, the mean denoised fMRI timeseries in each node was obtained by assigning FAAcoarse or FAAdetailed as an atlas file in CONN. The following analysis was performed using in-house software written for use with Matlab (2018b; The MathWorks).

### 2.7 Resting-state functional connectivity analysis using FAA

Functional connectivity (FC) was calculated as correlation coefficients (Pearson’s *r*) between BOLD-fMRI signals in each node. Actually, FAA enabled FC analysis across the anatomical hierarchy, such as FC between *inner* nodes, and FC between an inner node and leaf nodes, in addition to conventional FC between *leaf* nodes because FAA retains information related to anatomical hierarchy of brain structures (Standard FC analysis particularly addressing *leaf* nodes using the same MRI dataset is described in our earlier paper (Grandjean et al., 2020).). BOLD fMRI signals of an *inner* node were calculated as a size-weighted average of BOLD-fMRI signals in its descendant *leaf* nodes. Actually, FC analysis was restricted to nodes for brain parenchyma excluding nodes for fiber tracts and ventricles that were also defined in FAA. Statistical tests of correlation coefficients were applied as a mass univariate test using two-sided two-sample *t*-tests after the Fisher r-to-z transformation. Significance was set at a familywise false discovery rate (FDR) adjusted *p*-value of 0.05. Correlation coefficients for non-significant FC were set to 0 in Figs. 2d, 4a, and 4b. The inverse of Fisher transformation was applied to show correlation coefficients. Data are expressed as mean ± standard error of the mean (SEM) across animals, except where otherwise noted.

### 2.8 Data and code availability

Structural and functional MRI datasets used for this study are publicly available in the Brain Imaging Data Structure (BIDS) format at OpenNeuro (project ID: Mouse_rest_awake, https://openneuro.org/datasets/ds002551). Python codes for construction of FAA, and an HTML file using D3.js for zoomable visualization of anatomical hierarchy are available online (https://github.com/ntakata/flexible-annotation-atlas). The four FAAs are also available. Software environments were Python (3.7.1) using AllenSDK (version 0.16.1) written in Jupyter Notebook (5.6.0) on Anaconda (2018.12) on Windows 10 (Professional 64 bit, Microsoft). A yaml file for the anaconda environment is also available at the GitHub web page.

## 3. Results

### 3.1 Preprocessing of the original anatomical ontology data and annotation volume

AIBS provides a three-dimensional anatomical template (AT), annotation volume (AV), and a text file for anatomical ontology (AO) of the mouse brain (Fig. 1a). The original AV has 669 unique IDs for brain structures (nodes). The original AO defines an anatomical hierarchy of 1,327 nodes, among which 197 and 1,130 were inner nodes and leaf nodes, respectively (Materials and Methods section 2.2 gives definitions of inner nodes and leaf nodes). Combining or dividing nodes in the AV according to the anatomical hierarchy defined in the AO enabled creation of various AVs with arbitrary counts of nodes while maintaining the hierarchy. To achieve creation, it is crucially important to maintain consistency between AV and AO. Every leaf node and inner node in AO should and should *not* have corresponding voxels in AV, respectively. In this regard, *destructive* nodes exist in the original AO, which might impair the structure of the anatomical hierarchy in the AO and which might break consistency between AO and AV upon combining or dividing nodes.

Destructive nodes in the original AO can be classified as leaf nodes or inner nodes. Concretely, 490 *leaf* nodes in the AO have no corresponding voxels in the original AV. These *leaf* nodes are destructive because combining them in the AO to construct a new leaf node does not change the AV and thereby break one-on-one correspondence between the AO and AV. For instance, combining leaf node islands of Calleja (isl) and olfactory tubercle, molecular layer (OT1) to prepare a new leaf node olfactory tubercle (OT) in the original AO does not accompany modification of the original AV (Fig. 1b left). Similarly, 29 *inner* nodes in the original AO have corresponding voxels in the AV. They are also destructive because the size (voxel count) of these inner nodes changes depending on whether they are inner nodes or leaf nodes, resulting in node size inconsistency. For example, an *inner* node STR in the original AO has corresponding voxels of 1,392 nL in the original AV (Fig. 1b left). When the *inner* node STR becomes a *leaf* node by combining its child nodes such as CP and OT (Fig. 1b left), the size of the node STR changes to 22,457 nL, resulting in duplicated size assignment for the node depending on its node condition: either inner or leaf.

From preprocessing, destructive nodes in the original AO were eliminated, thereby creating a modified AO without destructive nodes, *i.e.* every leaf node and inner node in the modified AO respectively does and does not have corresponding voxels in the original AV. Specifically, destructive *leaf* nodes in the original AO were first eliminated, as were *leaf* nodes isl and OT1 in the AO (Fig. 1b). Then, destructive *inner* nodes in the AO were classified to two nodes: 1) an *inner* node without corresponding voxels in the original AV, and 2) a new *leaf* node with corresponding voxels in the AV (Fig. 1b). For example, an *inner* node STR in the original AO (Fig. 1b left) was divided to an inner node STR and a new leaf node STR_peri (Fig. 1b right).

Such a new *leaf* node was assigned a new, unique ID and was given a structure name and acronym of an original inner node, suffixed respectively with “_peripheral” and “_peri”. These suffixes were chosen because an original inner node was typically larger than a new *leaf* node. A mean size ratio of an original inner node over a new *leaf* node was 24 ± 8 for all 29 inner nodes.

This preprocessing caused modified AO, designated herein as AObase, and its corresponding AVbase. These AObase and AVbase were designated collectively as FAAbase, which had a total node count of 866, among which 197 and 669 were inner and leaf nodes, respectively. FAAbase is single-sided: homotopic areas on the right and left sides of the brain have the same ID and constitute a single node (Fig. 1a middle). Distribution by leaf node size in FAAbase revealed the existence of nodes that were too small or too large for fMRI analysis (blue symbols in Fig. 1c). Nodes smaller than a single voxel in fMRI (30 nL; dashed line in Fig. 1c) accounted for 6.7% of all leaf nodes in FAAbase (45 out of 669 leaf nodes). As one example, the node size of “Frontal pole, layer 6b” was only 2 nL (2 voxels in AVbase). By contrast, four nodes in the brain parenchyma were larger than 10,000 nL (dotted line in Fig. 1c; see Fig. 3a). An icicle plot depicts the anatomical hierarchy of brain structures in AObase (Fig. 1d).

### 3.2 Combining brain structures in AV while maintaining anatomical hierarchy

The proposed pipeline for FAA construction enables the combination of brain structures in AVbase while maintaining anatomical hierarchy in AObase (Fig. 2). To demonstrate this point, we combined nodes in FAAbase with a policy to make every node larger than a single voxel of fMRI by combining cortical layers and subregions in the brainstem and cerebellum, thereby obtaining FAAcoarse. For example, leaf node primary somatosensory area, nose, layer 1 (SSp-n1), layer 2/3 (SSp-n2/3), and others in the AObase (Fig. 2a left) were combined to create a new leaf node somatosensory areas (SS) in AOcoarse (Fig. 2a right). Consequently, FAAcoarse had a total node count of 101, among which 80 were leaf nodes. Comparison of icicle plots for FAAbase and FAAcoarse reveals that anatomical hierarchy was maintained during this node-combining process (Figs. 1d, 2b). FAAcoarse was constructed as double-sided, *i.e.* different IDs were assigned to homotopic nodes in the left and right side of the brain (Fig. 2c). The leaf node size distribution in FAAcoarse confirms that every node was larger than a single voxel of fMRI, although approximately 20% of leaf nodes were left extremely large (> 10,000 nL) (red symbols in Fig. 1c).

Resting-state FC across the anatomical hierarchy was calculated using FAAcoarse. Specifically, FC between inner nodes, and FC between a leaf node and an inner node were examined, in addition to conventional FC between leaf nodes (Fig. 2d). The FC matrix showed conspicuous vertical and horizontal red bands, suggesting that *inner* nodes, which are higher in the anatomical hierarchy in the brain, typically had higher FC in the cortex (Fig. 2d). One exception to this point was found in the prefrontal cortex (PFC; arrows in Fig. 2d). Whereas *leaf* nodes in the PFC such as the prelimbic area (PL), infralimbic area (ILA), and orbital area (ORB) had high FC within these *leaf* nodes (dense red region pointed by arrows in Fig. 2d), they showed low FC with inner nodes such as the isocortex and cortical plate (CTXpl) (pale red area indicated by arrowheads in Fig. 2d). This result apparently implies unique resting-state activity in the PFC, which was known to be vulnerable in psychiatric diseases (Chai et al., 2011; Sheline et al., 2010). A histogram of correlation coefficients in the FC matrix by FAAcoarse confirmed the tendency for higher FC in inner nodes (Fig. 2e, solid and dotted lines). Especially high correlation coefficients of approximately 0.8–1.0 were observed between inner nodes such as “Cerebrum and its child inner node Cerebral cortex” and “Cortical plate and its child inner node Isocortex” (Fig. 2e solid red line).

### 3.3 Dividing brain structures in AV objectively by gene expression or axonal projection data

Sizes of the largest four leaf nodes in the original AV exceed 10,000 nL (Fig. 3a). The pipeline for FAA construction enables classification of leaf nodes objectively using public resources by AIBS on gene expression and axonal projection. To demonstrate this point, the largest node field CA1 in the hippocampus (CA1) and caudoputamen (CP) were divided respectively based on gene expression and axonal projection to construct FAAdetailed (Fig. 3b).

A volume image of the gene-expression energy of a gene *Wfs-1* was retrieved from AIBS (Fig. 3c upper panels). *Wfs-1* is expressed densely at dorsal part in the hippocampal CA1 region when compared to its ventral part (Cembrowski and Spruston, 2019). A thresholding histogram of the gene-expression energy in a leaf node CA1 (Fig. 3c right histogram) indicated the node as divisible into two: nodes with lower or higher gene expression “CA1_geneL” or “CA1_geneH”, respectively (blue and red areas in Fig. 3c bottom panels). Similarly, volume images for density of axonal projection from the agranular insula area (AI) in the cortex to CP in the right hemisphere were retrieved from AIBS (Fig. 3d upper panels). Ventral CP receives cortical afferents mainly from the insular cortex (Berendse et al., 1992; Steiner and Van Waes, 2013). According to a thresholding histogram of the axonal projection density in a leaf node CP (Fig. 3d right histogram), the nodes were classifiable into two types: nodes with lower and higher projection density “CP_fiberL” and “CP_fiberH”, respectively (blue and red areas in Fig. 3d bottom panels). More specifically, 25 experimental data for axonal projection density from AI in the right cortex to CP were retrieved from AIBS (June 6, 2019). Representative projection data are depicted in Fig. 3d upper panels. The max projection of these 25 volumes was flipped horizontally to obtain symmetric data for dividing the node CP (Fig. 3d lower panels).

This procedure of division modified AObase, which we designated as AOdetailed, and yielded its corresponding annotation volume, AVdetailed. These were collectively designated as FAAdetailed, which had a total node count of 1,381, among which 1097 and 284 were leaf nodes and inner nodes, respectively (Fig. 3e). The divided nodes were visible in AVdetailed (arrows and arrowheads in Fig. 3f). Distribution of the leaf node size in FAAdetailed indicates that the largest nodes CA1 and CP in the original FAAbase were removed in FAAdetailed (deep green plots in Fig. 1c).

Resting-state FC was calculated using FAAdetailed (Fig. 4a). The divided nodes in CA1 or CP showed distinct FCs in their strength and patterns. Specifically, there were 229 significant FCs from a node CA1_geneL (ventral CA1) and/or a node CA1_geneH (dorsal CA1) in the FC matrix (*n* = 13, paired *t*-tests with FDR adjusted *p*-value of 0.05). Among the FCs, 88.7% (203 among 229 FCs) were significantly larger for the ventral CA1 than the dorsal one (*n* = 13, paired *t*-test with FDR adjusted *p*-value of 0.05; blue markers at the top of Fig. 4b upper panel), suggesting stronger FCs from the ventral CA1 than from the dorsal CA1. Furthermore, 80.8% of the FCs (185 among 229 FCs) were significant only for either ventral or dorsal CA1, implying distinct patterns of FCs from those of ventral and dorsal CA1. Similarly, there were 196 significant FCs from a node CP_fiberH (ventral CP) and/or a node CP_fiberL (dorsal CP) in the FC matrix (*n* = 13, paired *t*-tests with FDR adjusted *p*-value of 0.05). Among the FCs, 97.5% (191 among 196 FCs) were significantly larger for the ventral CP than the dorsal CP (*n* = 13, paired *t*-test with FDR adjusted *p*-value of 0.05; red markers at the top of Fig. 4b lower panel), suggesting stronger FCs from the ventral CP than that from the dorsal CP. Furthermore, more than half of the FCs (59.2%; 116 among 196 FCs) were significant only for either ventral or dorsal CP, implying distinct patterns of FCs from those of ventral and dorsal CP.

### 3.4 Resting-state functional connectivity across the anatomical hierarchy and between cortical layers

Earlier investigations of hierarchical organization of resting-state FC have specifically examined *leaf* nodes (Doucet et al., 2011; Gotts et al., 2020); Consequently, modulation of FC across the anatomical hierarchy has been little investigated because examination of *inner* nodes was necessary. As an example of FC modulation across the hierarchy, one can presume a situation in which FC from a source leaf node S to an *inner* node A is higher than that to *leaf* node B or C, which are descendants of the inner node A (Fig. 4c). This situation suggests a mechanism to modulate FC in larger units in the brain, *i.e.* an *inner* node might be able to constitute a larger unit to control the activities of its descendant leaf nodes.

To investigate the case, an FC matrix obtained with FAAdetailed was examined in a search for source *leaf* nodes S that met the following three criteria: 1) FC between S and an *inner* node A was larger than 0.8; 2) the inner node A was not an ancestor of the leaf node S, and 3) FC between them was larger than FC from node S to the leaf nodes (B and C) which were descendants of inner node A. FAAdetailed had 1,363 grey-matter nodes, among which 277 were inner nodes and 1,086 were leaf nodes. Among the leaf nodes, 31% (339 leaf nodes) passed the first criterion. Among the nodes, 5% (16 leaf nodes) passed the second criterion. Furthermore, finally, 81% (13 leaf nodes) passed the last criterion. Diamonds in Fig. 4a denote pairs of a leaf node and an inner node that satisfied the criteria.

To visualize the FC distribution explicitly with anatomical hierarchy, FCs from one of the 13 leaf nodes, “Retrosplenial area, ventral part, layer 2/3_L” (RSPv2/3_L; node size, 518 nL), were represented as CoSE representation in Fig. 4d. FC from the source *leaf* node RSPv2/3_L (yellow start in Fig. 4d) to an *inner* node “Retrosplenial area, ventral part_R” (RSPv_R, 2241 nL; correlation coefficient, 0.82 ± 0.01; *n* = 13; arrow in Fig. 4d) was indeed larger than FC from the source node to descendant leaf nodes of the inner nodes (arrowheads in Fig. 4d), *i.e.* RSPv1_R (517 nL, 0.80 ± 0.02), RSPv2/3_R (549 nL, 0.81 ± 0.02), RSPv5_R (828 nL, 0.73 ± 0.03), RSPv6a_R (326 nL, 0.48 ± 0.04), and RSPv6b_R (21 nL, 0.39 ± 0.04). Statistical tests revealed, however, that FC from the source node to the *inner* node was not always significantly larger than FCs to the descendant *leaf* nodes. Actually, FC from the source node to the inner node and FCs to the descendant leaf nodes (RSPv1_R and RSPv2/3_R) were not significantly different (*n* = 13, repeated measures ANOVA with post-hoc test using Holm-adjusted *P* value of 0.05). Similar results were obtained for all 13 leaf nodes.

Cortical-layer-specific FC analysis was attempted using FAAdetailed because thin but large nodes for cortical layers were defined in it. Marked variation of FC within cortical layers was observed in most cortical regions. For example, in the primary somatosensory area (barrel field in the left cortex; SSp-bfd), supragranular (L1 and 2/3) and infragranular (L5, 6a, and 6b) layers showed higher FC within them, but lower FC between them (Fig. 4e). Specifically, FC between L1 and L2/3 (correlation coefficient, 0.64 ± 0.07; *n* = 13) and FC between L5 and L6a (0.85 ± 0.02) were, respectively, significantly larger than FC between L2/3 and L6a (0.28 ± 0.03) (*n* = 13, one-way ANOVA with post-hoc Tukey-HSD test using a *P*-value of 0.05). Negative FCs (−0.16 ± 0.04) between supragranular layer (L1) and infragranular layer (L6b) were disregarded because L6b was small (77 nL) and because it might not satisfy the assumptions presented below.

## 4. Discussion

The absence of a *de facto* standard annotation atlas for the mouse brain in MRI studies might engender inconsistent ROI definitions for brain structures, making it difficult to compare, replicate, and validate results obtained from other laboratories (Pallast et al., 2019). In fact, multicenter comparison of resting-state fMRI on the mouse brain has not been reported until this year (Grandjean et al., 2020). This situation conflicts with a recent trend for data-sharing in neuroscience (Ascoli et al., 2017; “Data sharing and the future of science,” 2018; Nichols et al., 2017). The current study addressed this issue by constructing a pipeline for a flexible annotation atlas (FAA). Actually, FAA presents three advantages. The first is flexibility: FAA enables preparation of an annotation atlas with a total count of brain structures from 1 to over 1,000, while maintaining anatomical hierarchy. Considering the huge amount of public resources provided by AIBS related to gene expression and axonal projection, the proposed FAA pipeline can prepare innumerable patterns of an annotation atlas that would meet the various needs of researchers. The second is simplicity: Construction of FAA requires only two steps. Manual editing of a text file and running a Python code with user-specified text-based information after preprocessing. Third is reproducibility: Combination and division of nodes were performed objectively by the codes using text-based information. Consequently, sharing of whole brain ROI definition among laboratories would be facilitated through text-based information.

FAA enabled FC analysis including *inner* nodes in addition to leaf nodes, thereby providing an opportunity to examine FC across the anatomical hierarchy. The uniqueness of resting-state activity in the PFC was inferred from results of FC analysis with FAAcoarse. Furthermore, modulation of FC by larger units in the brain, *i.e.* an inner node, was examined using FAAdetailed. Whereas some pairs of a source leaf node and an inner node indeed had higher FC than pairs of the source node and descendant leaf nodes of the inner node, the difference of FCs was not always significant, and not supporting the idea of FC modulation by an inner node, at least during resting-state *static* FC of awake mice (Grandjean et al., 2017a). Therefore, a more likely interpretation for higher FC between a source leaf node and an inner node observed here might be an SNR increase by averaging of BOLD-fMRI signals in descendant leaf nodes of the inner node. Irrespective of the negative result, FC across the anatomical hierarchy is worthy of further investigation during, say, a situation with large-scale brain dynamics such as cortical spreading depression (Yoshida et al., 2015). Furthermore, only *leaf* nodes were examined as source nodes in this study. Consequently, the possibility of inner nodes as source nodes was not explored.

Laminar fMRI is an emerging field for deciphering interaction of the brain, discriminating input layers and output layers in the cortex (Goense et al., 2012; Norris and Polimeni, 2019; Scheeringa and Fries, 2017; Yu et al., 2014). Resting-state FC analysis between cortical layers with fMRI is under active investigation with challenges related to spatial resolution of fMRI and vascular distribution in the cortex (Mishra et al., 2019; Poplawsky et al., 2019). In the present study, cortical layer-specific FC analysis was attempted using FAAdetailed, which defined thin but large nodes for cortical layers. This analysis assumed that BOLD-signal fluctuation was uniform within each cortical layer, and that contaminating signals in a node from other nodes were cancelled out by averaging signals within the target node. For example, the cortical layer thickness in SSp-bfd is represented by only 1–5 voxels (100–500 μm) in an annotation atlas, FAAdetailed, that was comparable to or even smaller than the in-plane resolution of a single voxel in fMRI (200 × 200 μm^2^). Still, node for each cortical layer was 13–25 times larger than a single voxel of fMRI (30 nL), except for layer 6b (77 nL). Significant variation in FC within *vs.* between supra-granular and infragranular layers was found in most cortical regions, which is apparently consistent with results of earlier studies examining laminar differences of neuronal activity measured using calcium imaging (Ayaz et al., 2019), electrophysiology (Sakata and Harris, 2009), and fMRI (Mishra et al., 2019). It is noteworthy that such layer-specific FC was observed both in the dorsal (*e.g.* Primary somatosensory area) and lateral (*e.g.* Agranular insular area) parts of the cortex, implying that artifacts caused by spatial registration of fMRI images did not account for the variation considering the fact that the Allen reference atlas was slightly expanded dorsoventrally, but not mediolaterally. Further investigation must be conducted to assess, interpret, and validate laminar FC analysis of fMRI data using thin but large nodes in FAA.

One limitation of the proposed FAA is that combining a part of sibling nodes is not allowed in FAA construction. All descendant nodes of an inner node should be combined to make the inner node a new leaf node. This all-or-none rule for combining descendant nodes was necessary to maintain consistency of anatomical hierarchy in FAA. Even overly small nodes were reserved if their large sibling nodes were intended for use in additional analysis. For example, in construction of FAAdetailed, a node for layer 6b (1 nL) was not combined with other nodes to reserve its sibling nodes in the cortical frontal pole such as layer 1 (119 nL), layer 2/3 (122 nL), and layer 5 (186 nL). Consequently, 188 nodes (17.1% of all leaf node) in FAAdetailed were smaller than a single voxel of fMRI. However, the volume occupied by these small nodes was negligible: 0.6% of the whole brain volume.

Application of FAA is not limited to MRI data. Whole-brain imaging study with tissue-clearing techniques or a serial-sectioning imaging system would benefit from consistent and flexible ROIs provided by FAA, once registered to the Allen reference atlas (Richardson and Lichtman, 2015; Seiriki et al., 2017; Susaki et al., 2014). Two-dimensional wide-field cortical imaging can also use FAA (Barson et al., 2020; Fan et al., 2019; Matsui et al., 2016; Sofroniew et al., 2016) through a flat-map projection of the Allen reference atlas (Harris et al., 2019; Weed et al., 2019). FAA facilitates integration of these data with open source resources for which datasets are registered to the Allen Brain Atlas (EPFL, 2005; Janelia Research Campus, 2017), thereby promoting cellular level whole-brain investigation (Fürth et al., 2018).

## Acknowledgements

The authors would like to thank Drs. Yoshifumi Abe and Joanes Grandjean for critical comments on the manuscript.

## Funding

This research was supported by AMED under Grant Number JP19dm0207069 and by JSPS KAKENHI Grant Numbers 16K07032, 16H01620, 18H04952.

## Supplementary Figures

**FigS 1.**
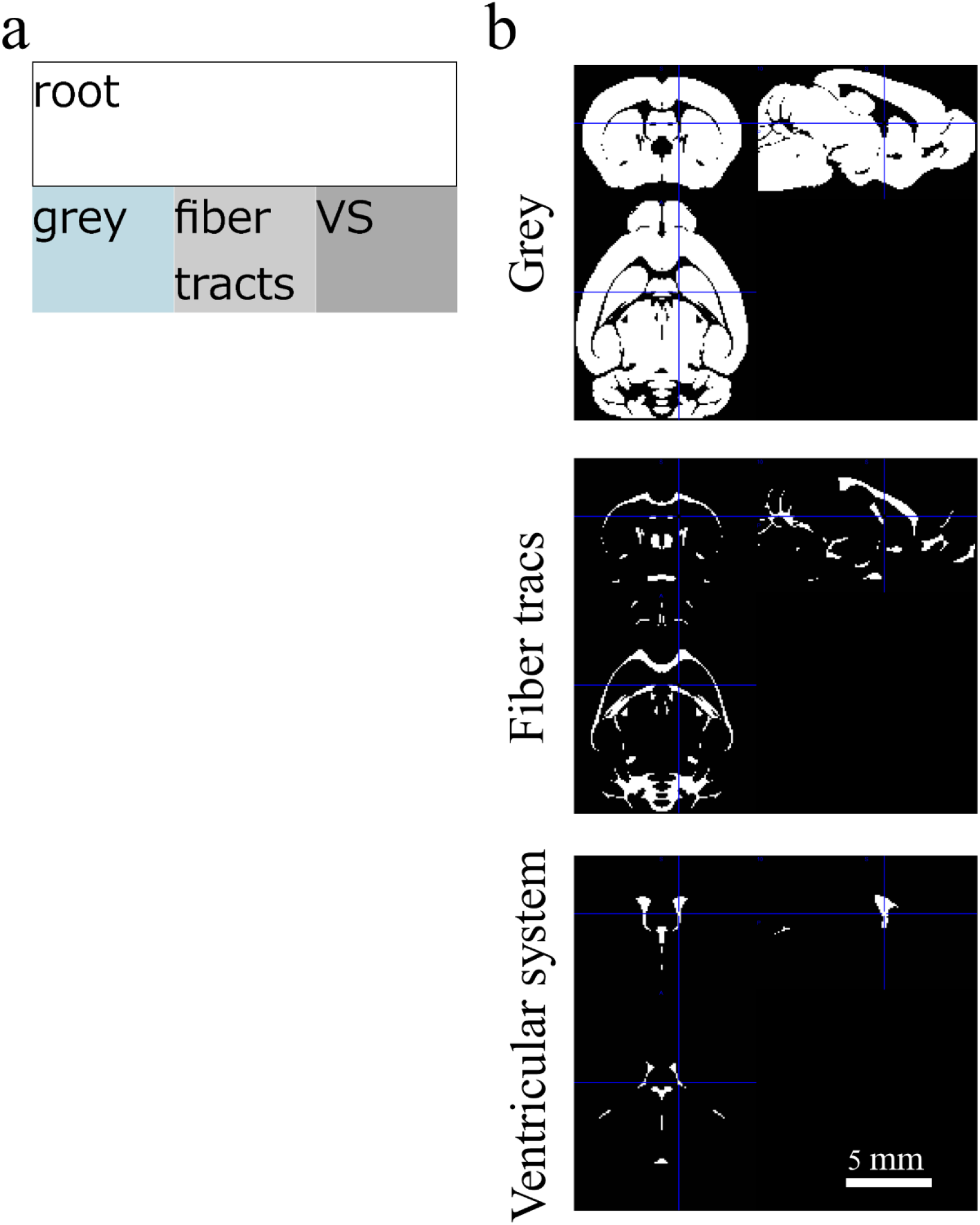
FAAsegment for brain tissue segmentation. **(a)** An icicle plot showing the anatomical hierarchy of FAAsegment that has four nodes in total, of which three are leaf nodes corresponding to grey matter, fiber tracts, and ventricular system. **(b)** Coronal, sagittal, and horizontal planes of three leaf nodes in FAAsegment that was used for brain tissue segmentation during temporal preprocessing of fMRI data.

**FigS 2.**
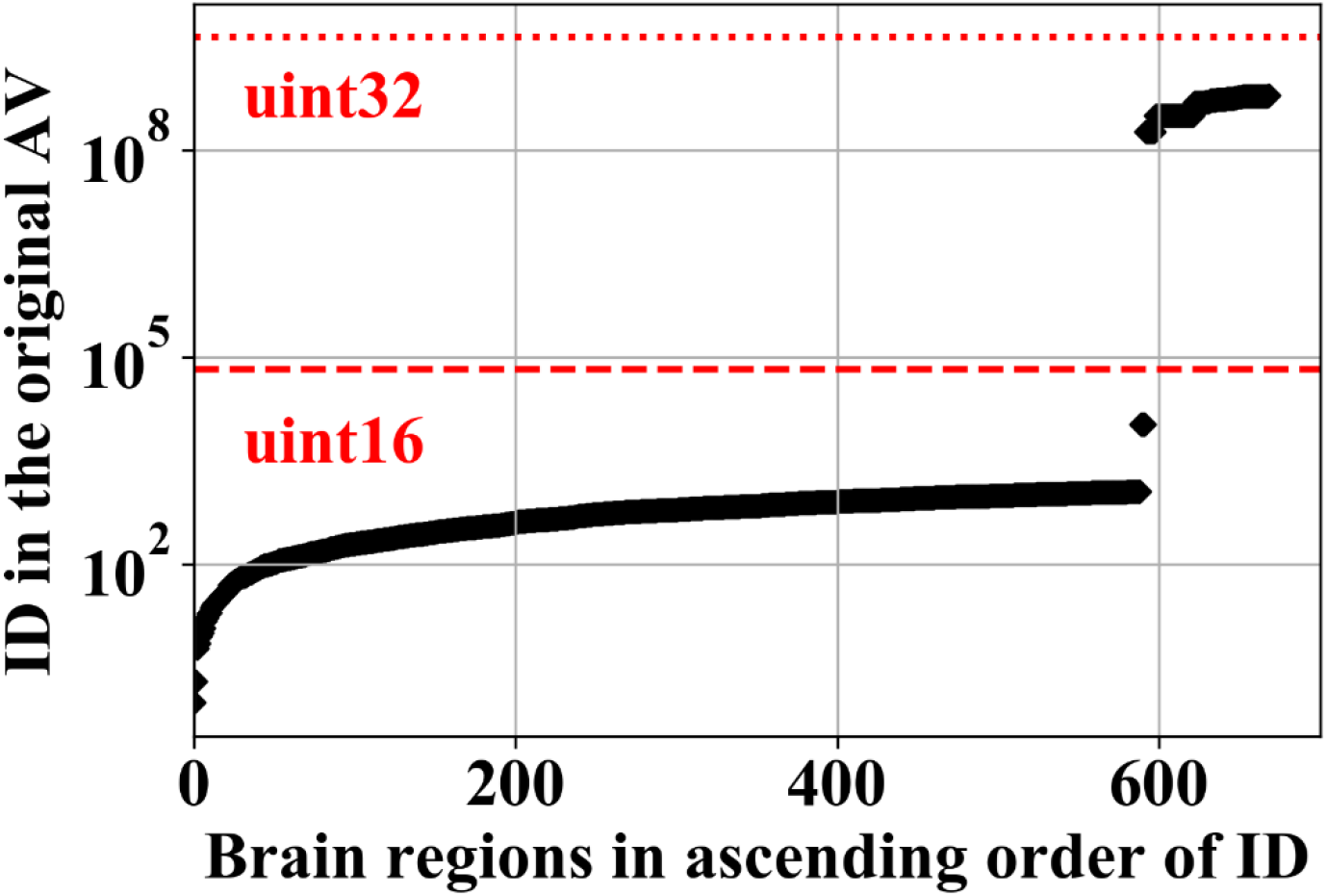
IDs for brain structures in the original AV. The original AV is 32-bit UINT with 670 unique integer values that correspond to an ID for a brain structure. More than 11% of IDs (77 IDs out of 670) exceeded the maximum value for 16-bit UINT (2^16^ = 65,536, a red dashed line just below 10^5^). Remapping of IDs in the original AV and AO was implemented in a construction pipeline for FAA to put them within the range of 16-bit UINT because some MRI viewers such as ITK-SNAP (www.itksnap.org/pmwiki/pmwiki.php) and Mango (ric.uthscsa.edu/mango/) did not support 32-bit UINT. Red dotted line: The maximum value for 32-bit UINT (2^32^ = 4,294,967,296).

## Supplementary Tables

**Supplementary Table 1.** Download links for online resources by Allen Institute for Brain Science.

| File name/contents | Description |
| --- | --- |
| average_template_25.nrrd,<br>average_template_50.nrrd,<br>average_template_100.nrrd | Average template (AT) of the mouse brain in NRRD 16-bit UINT format (http://teem.sourceforge.net/nrrd/) with spatial resolution of 25, 50, and 100 $\mu\text{m}$ isovoxel. Volume dimensions are $528 \times 320 \times 456$, $264 \times 160 \times 228$, and $132 \times 80 \times 114$ in PIR (+x = posterior, +y = -inferior, +z = right) coordinate, respectively, with an image-origin at anterior upper left corner long unit in micrometers.<br>DL: http://download.alleninstitute.org/informatics-archive/current-release/mouse_ccf/average_template/<br>Ref: http://help.brain-map.org/download/attachments/2818169/MouseCCF.pdf?version=1&modificationDate=1432933021016&api=v2 |
| annotation_10.nrrd,<br>annotation_25.nrrd,<br>annotation_50.nrrd,<br>annotation_100.nrrd | Annotation volume (AV) of the mouse brain in NRRD 32-bit UINT format with spatial resolution of 10, 25, 50, and 100 $\mu\text{m}$ isovoxel. Volume dimensions are $1320 \times 800 \times 1140$, $528 \times 320 \times 456$, $264 \times 160 \times 228$, and $132 \times 80 \times 114$ in PIR coordinates. AVs with higher spatial resolution include additional brain structures [acronym (ID) for a brain structure]: c (10), tspd (1), and RSPd4 (2).<br>DL: http://download.alleninstitute.org/informatics-archive/current-release/mouse_ccf/annotation/ccf_2017/<br>Ref: http://help.brain-map.org/display/mouseconnectivity/API |
| 1.json | A text file in JSON format that defines annotation ontology (AO) of the mouse brain (Structure graph ID: 1).<br>DL: http://api.brain-map.org/api/v2/structure_graph_download/1.json<br>Ref: http://help.brain-map.org/display/api/Atlas+Drawings+and+Ontologies |
| Gene expression energy | Volume data for gene expression energy for approximately 20,000 genes in the adult-mouse brain in MetaImage 32-bit FLOAT format (https://github.com/Kitware/MetaIO) with spatial resolution of 200 $\mu\text{m}$ isovoxel. Their respective volume dimensions are $67 \times 41 \times 58$ in PIR coordinates.<br>Ref: http://help.brain-map.org/display/mousebrain/API |
| Axonal projection density | Volume data for axonal projection density in the brain in NRRD 32-bit FLOAT format with spatial resolutions of 10, 25, 50, and 100 $\mu\text{m}$ isovoxel. Their respective volume dimensions are $1320 \times 800 \times 1140$, $528 \times 320 \times 456$, $264 \times 160 \times 228$, and $132 \times 80 \times 114$ in PIR coordinates.<br>Ref: http://help.brain-map.org/display/mouseconnectivity/API |

DL, download link; Ref, reference site. Data were retrieved June 12, 2019.

**Supplementary Table 2.** Steps to construct FAA of the mouse brain.

| Step | Code/procedure | Input | Primary output |
| --- | --- | --- | --- |
| 0 | Prepare_AObaseAVbase.ipynb | Original resources at AIBS<br>- 1.json (anatomical ontology text-file)<br>- annotation_100.nrrd (annotation volume)<br>This “preprocessing” eliminates *destructive* brain structures (nodes) in the original resources, performed by a Python code without user inputs. | FAAbase comprises<br>- AObase.json<br>- AVbase.nrrd |
| 1 | Combine brain structures | - AObase.json<br>This first step combines leaf-nodes to obtain a new leaf-node with larger volume while maintaining anatomical hierarchy by manually editing a JSON-formatted text file, AObase.json. Specifically, copy AObase.json and rename it to AObase_c.json. Then, to combine all descendent nodes of an *inner* node, delete all contents within brackets [] of a key “children” for the inner node. This would be facilitated by a text editor such as Vim (https://www.vim.org/) with functionality to jump to matching brackets. To support this further, a zoomable plot of an anatomical hierarchy in AObase_c.json is provided as an HTML file using D3.js (https://d3js.org/) (e.g. Fig. 1d). To open this HTML-file, it is recommend that “Web Server for Chrome” (github.com/kzahel/web-server-chrome) be used. It is available at the Chrome web store because direct access to a local file is prohibited for security reasons in a recent web-browser, e.g. Firefox after ver. 68.0. | - AObase_c.json |
| 2 | Divide_nodes.ipynb | A user-specified text-based information on<br>- **AObase_c.json**<br>- **Target_ROI_IDs** (brain structures) defined in AObase_c.json for dividing nodes<br>- **ExpID** defined at AIBS to specify a gene of interest<br>- **Acronyms** for brain structures to specify source- and target-node for axonal projection<br>This second step divides leaf nodes based on gene expression and axonal fiber projection using a Python code with a user-specified text-based information, resulting in a flexible annotation atlas (FAA) that comprises an annotation ontology (AO) text-file and an annotation volume (AV). Five additional modifications are performed during this step: 1) assigning different IDs for homotopic nodes in the right and left side of the brain to make annotation atlas bilateral, e.g. Fig. 2c; 2) remapping IDs for brain structures in the original AO and AV to be in the range of 16-bit UINT; 3) transforming a format of the original AV from NRRD to NIfTI-1 (https://nifti.nimh.nih.gov/) because some programs such as MRICron and SPM do not support NRRD format; 4) modifying image orientation from posterior-inferior-right (PIR) to right-anterior-superior (RAS) as is widely used in NIfTI standard; and 5) setting spatial origin of AV and AT to the bregma referring to the mouse brain atlas (Paxinos and Franklin, 2001), from that at top left corner of a volume image. This step enables specification of the location of the brain structure using mm-coordinates, as in human MRI study. There are 6 nodes located only in the midline: Medulla, behavioral state related (MY-sat), its child nodes (nucleus raphe magnus (RM), pallidus (RPA), and obscurus (RO)), vascular organ of the lamina terminalis (OV), and Edinger-Westphal nucleus (EW). These nodes were assigned to the right side of the brain during the first step. For reconstruction of FAA, share the text-based information shown in bold face above: AObase_c.json, Target_ROI_IDs, ExpID, and Acronyms (Fiber_from and Fiber_to). | **FAA** comprises<br>- **AO_LR_remapID.json**<br>- **AV_LR_remapID_RAS.nii** |

**Supplementary Table 3.** Subfunctions in Prepare_AObaseAVbase.ipynb for preprocessing.

| Step | Code | Input | Output |
| --- | --- | --- | --- |
| 1 | Add_VC_to_AO.ipynb | - 1.json<br>- annotation_100.nrrd | - 1_VC.json |
|  | A voxel count (VC) information for a brain structure in AV is appended to an anatomical ontology file. |  |  |
| 2 | Prune_leaf_ROI_wo_VC_in_AO.ipynb | - 1_VC.json | - 1_VC_pruned.json |
|  | Destructive leaf nodes (leaf nodes without voxel counts in AV) are deleted from AO. |  |  |
| 3 | Divide_internal_ROI_with_VC_in_AO.ipynb | - 1_VC_pruned.json | - 1_VC_pruned_divided.json<br>- dividedIDs.csv |
|  | Destructive inner nodes (inner nodes with voxel counts in AV) are divided, leading to a new leaf-node with new ID. Its name and acronym are the same as the original ones suffixed respectively with “_peripheral” and “_peri”. |  |  |
| 4 | Update_ID_in_AV_to_reflect_divided_AO.ipynb | - annotation_100.nrrd<br>- dividedIDs.csv | - AVbase.nrrd |
|  | IDs in AV is updated according to an anatomical ontology file, 1_VC_pruned_divided.json. |  |  |
| 5 | Add_VC_to_AO.ipynb | - AVbase.nrrd<br>- 1_VC_pruned_divided.json | - AObase.json |
|  | Update voxel-count information in the AO according to AVbase.nrrd. |  |  |

**Supplementary Table 4.**
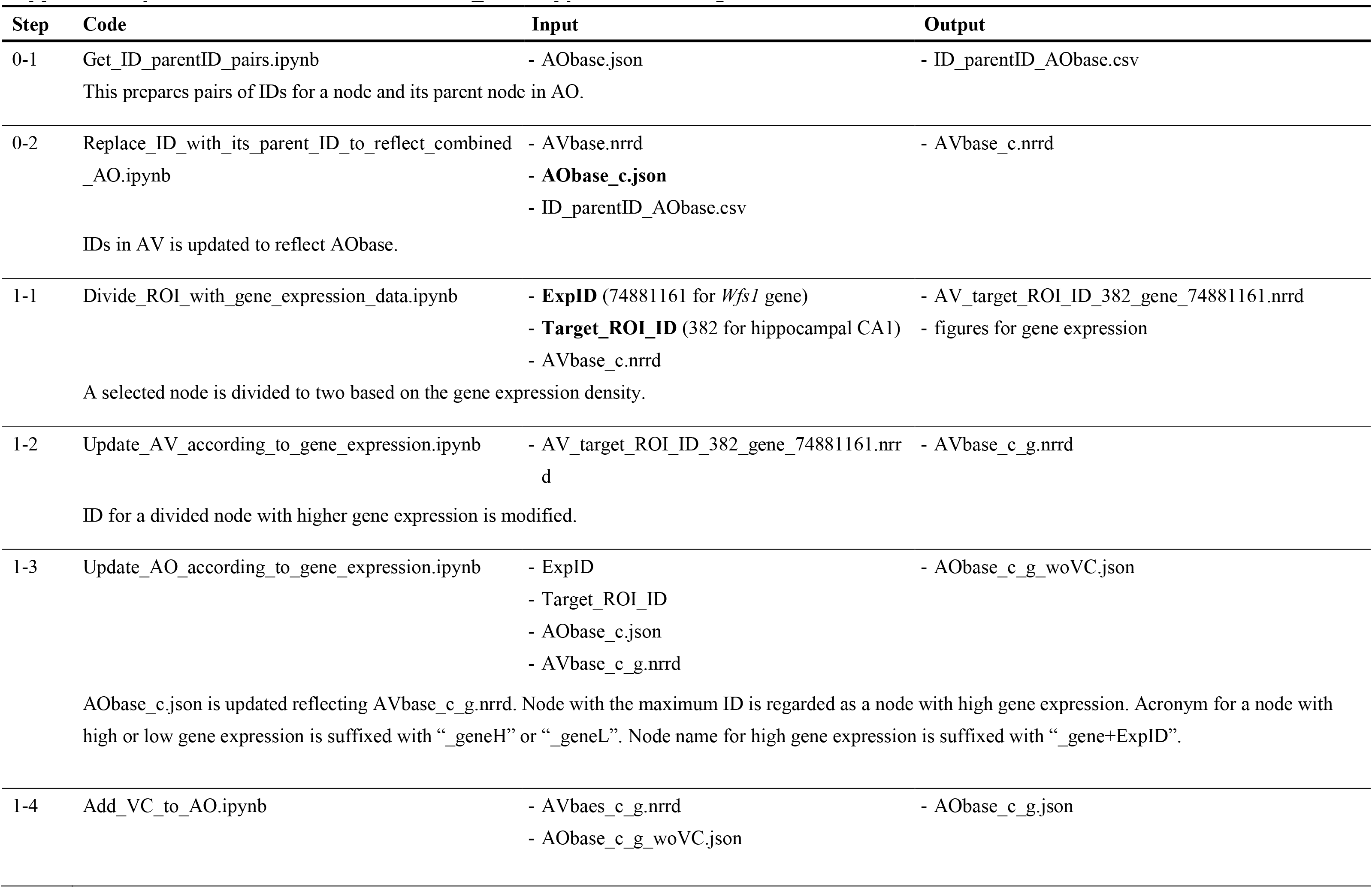
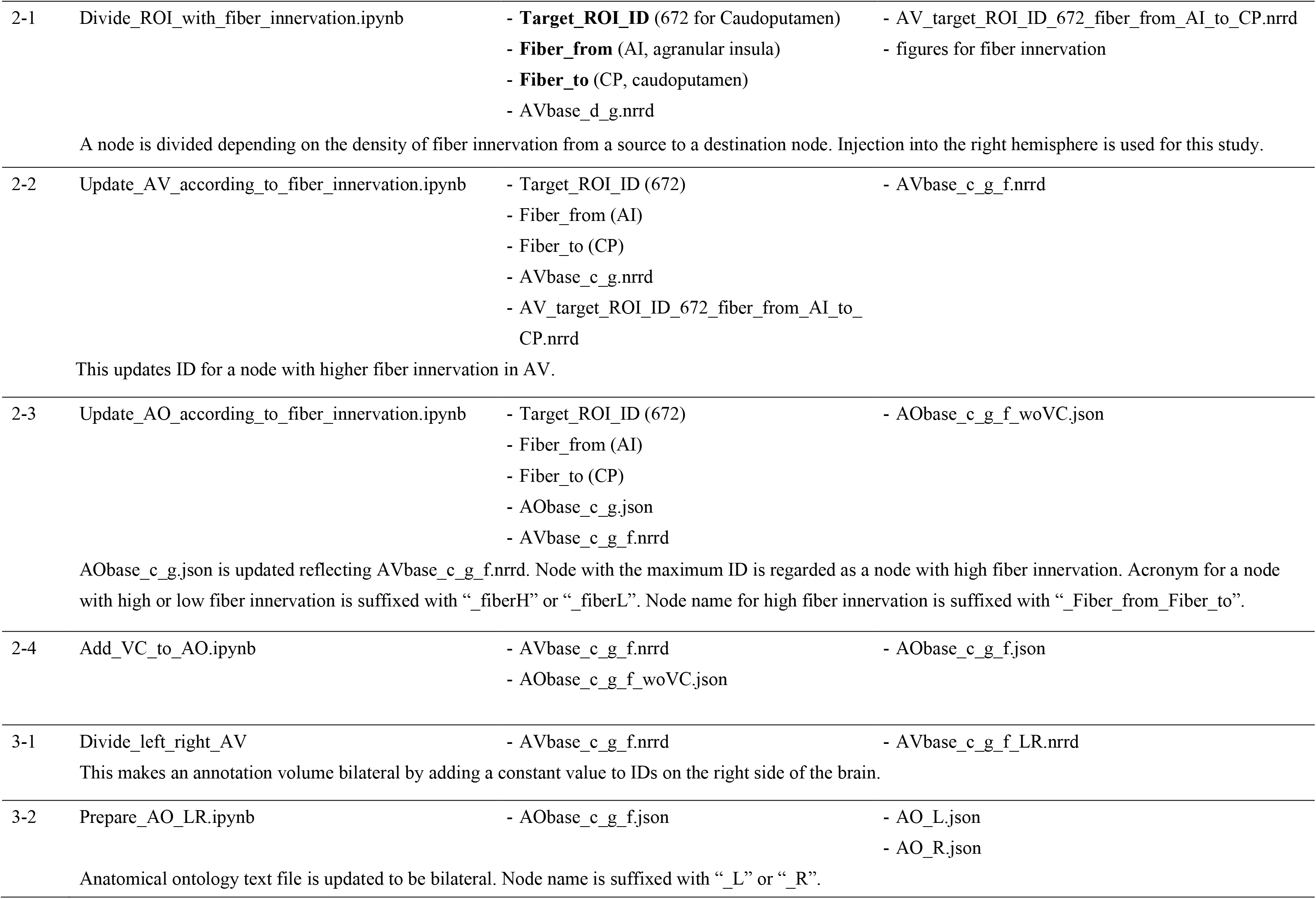
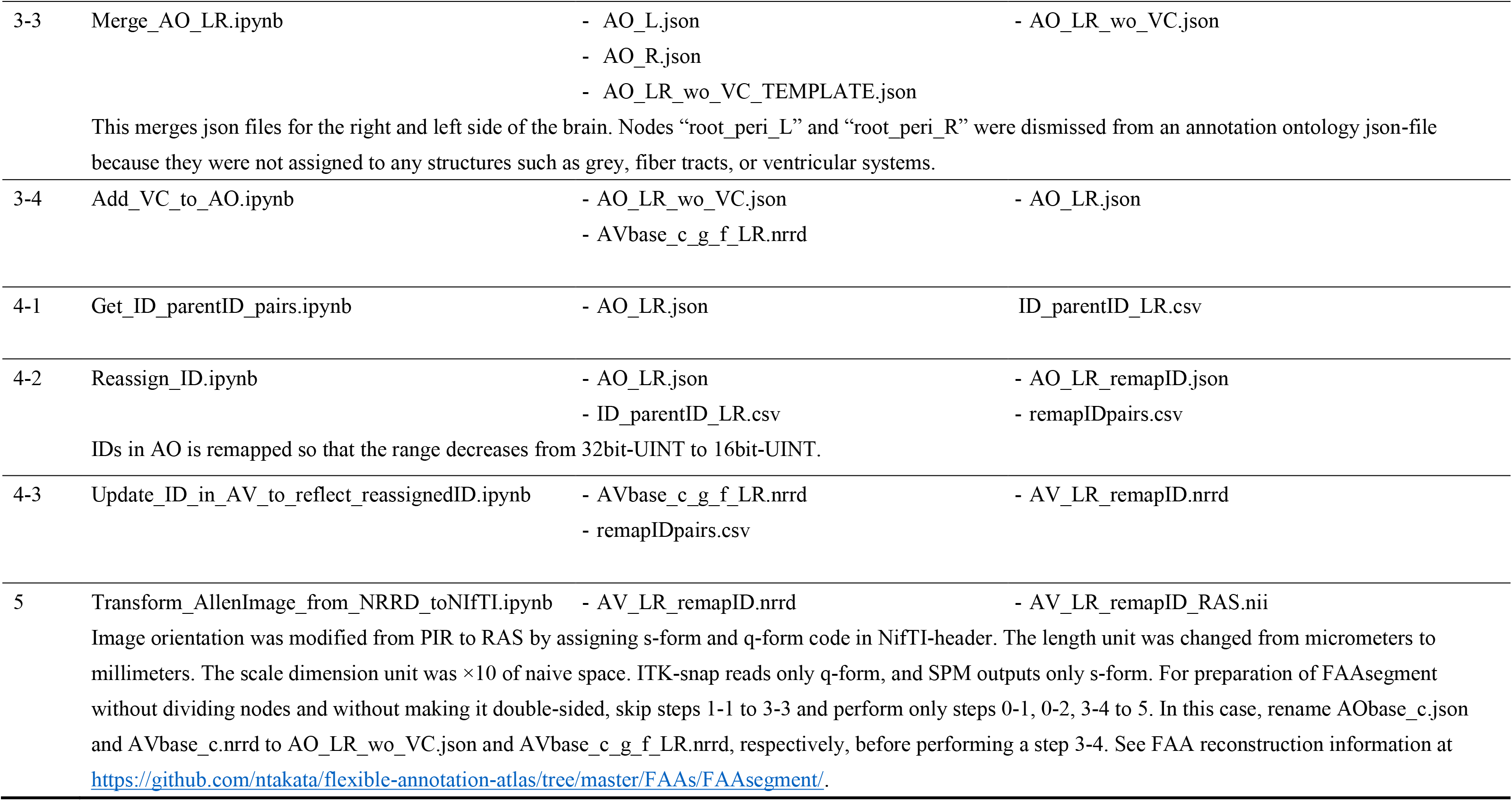
Subfunctions in Divide_nodes.ipynb for dividing leaf-nodes.

## Notes

#### Summary of Updates

Supplementary Tables added; Three additional references; Funding information updated.

https://github.com/ntakata/flexible-annotation-atlas

https://openneuro.org/datasets/ds002551

